# Parasitic wasps do not lack lipogenesis

**DOI:** 10.1101/2021.03.16.435583

**Authors:** Joachim Ruther, Lorena Prager, Tamara Pokorny

## Abstract

Fatty acids are crucial primary metabolites for virtually any creature on earth. Therefore, most organisms do not rely exclusively on nutritional supply with fatty acids but have the ability to synthesize fatty acids and triacylglycerides de novo from carbohydrates, a process called lipogenesis. The ubiquity of lipogenesis has been questioned by a series of studies reporting that many parasitic wasps (parasitoids) do not accumulate lipid mass despite having unlimited access to sugar. This has been interpreted as an evolutionary metabolic trait loss in parasitoids. Here, we demonstrate de novo biosynthesis of fatty acids from ^13^C-labeled α-D-glucose in eleven species of parasitoids from six families. We furthermore show with the model organism *Nasonia vitripennis* that lipogenesis occurs even when lipid reserves are still intact, but relative ^13^C-incorporation rates increase in females with widely depleted fat reserves. Therefore, we conclude that the presumed “lack of lipogenesis” in parasitoids needs to be re-evaluated.

## Introduction

Fatty acids and their derivates are crucial primary metabolites of virtually any organism on earth. They fulfil indispensable functions such as forming biomembranes of cells and cell organelles, storing metabolic energy, or being precursors of hormones, pheromones, and other signalling molecules [1–3].

This leads to a permanent demand for fatty acids which is not only covered by nutritional supply but also by the ability to synthesize fatty acids de novo from other primary metabolites such as carbohydrates. Fatty acid biosynthesis from sugar-derived acetyl coenzyme A units and subsequent incorporation of the fatty acids into triacylglycerides (TAGs) is commonly referred to as lipogenesis [4].

Some authors, however, use the term lipogenesis less restrictively for the de novo synthesis of fatty acids from carbohydrates [5, 6] irrespective of the purpose these fatty acids are used for. Given the crucial importance of fatty acids, lipogenesis is highly conserved in organisms and the vast majority of them possess the enzymes necessary for this basic metabolic pathway [7].

In insects, the ubiquity of lipogenesis has been questioned in parasitoid wasps [8, 9], one of the most speciose groups of insects on earth [10], that develop parasitically in or on other arthropod hosts [11, 12]. Several papers have reported experiments in which parasitoid wasps and other insect species with a parasitic lifestyle, despite having unlimited access to sugar sources, did not gain lipid mass [8, 9, 13–19]. The lack of lipid mass gain in feeding experiments as well as supporting gene expression data [19] have been interpreted as a general inability of parasitoid wasps to synthesize fatty acids and storage lipids such as TAGs from sugars. This “lack of lipogenesis” in parasitoid wasps has been suggested to be an example of an evolutionary metabolic trait loss enabled by environmental compensation [9]. It has been argued that parasitoid wasps are provided with sufficient amounts of lipids by their hosts, making the ability to synthesize fatty acid derivatives redundant. The ability of some parasitoid species to manipulate their host’s metabolism in that these produce or release higher amounts of lipids seemed to support this hypothesis [20–23]. However, even if this mechanism were a general feature in parasitoid-host interactions, it would merely benefit the parasitoids’ juvenile stages and not rule out that adults replenish ebbing lipid resources through lipogenesis. Apart from this, studies claiming a general lack of lipogenesis in parasitoid wasps suffer from some conceptional and methodical flaws that challenge the generality of the hypothesis. First, in some of these studies not all studied species failed to gain lipid mass upon sugar feeding [9, 24]. These examples, however, have not been used to question the generality of the rule but rather considered as rare exceptions in which lipogenesis has re-evolved [9]. Secondly, many studies investigating lipogenesis in parasitoids applied relatively unselective bulk methods for lipid quantification such as gravimetry [9, 16, 17] or colorimetry [13–15, 18, 25]. These approaches, however, do not allow for a discrimination between fatty acid derivatives and other lipids. Furthermore, while being suitable to monitor mass gains and losses of lipids, gravimetric and colorimetric methods are not suited to conclude the complete lack of a basic metabolic pathway such as lipogenesis. While constantly losing lipid mass throughout their lifetime, parasitoids might very well use encountered carbohydrate sources such as floral and extrafloral nectar or honeydew [26, 27] to replenish depleting lipid resources. This would remain unnoticed by bulk techniques if lipogenesis and fatty acid degradation occurred simultaneously. Without performing suitable labelling experiments and applying high-resolution analytical techniques such as gas chromatography/mass spectrometry (GC/MS) to scrutinize the extracted fatty acid derivatives for the presence of sugar-derived atoms, the lack of lipogenesis in parasitoid wasps cannot be claimed. This view has been supported by a recent study [28] on *Nasonia vitripennis* (Nv), a model organism for the study of parasitoid wasp biology [29], for which several studies have claimed a lack of lipogenesis [8, 9, 19, 30]. Feeding experiments involving fully ^13^C-labeld α-(+)-D-glucose conducted with Nv and three other species of the so-called *Nasonia* group [31], i.e. *N. giraulti*, *N. longicornis*, and *Urolepis rufipes*, unequivocally demonstrated the incorporation of sugar-derived carbon into fatty acids as well as into the male sex pheromone which is synthesized from fatty acid precursors [32–34]. Furthermore, a proteomics approach revealed that several enzymes involved in fatty acid biosynthesis such as acetyl-CoA carboxylase, fatty acid synthase, and ATP-citrate synthase are highly abundant in the male pheromone gland [28]. A rapid decrease of lipids has been found in Nv during the first 72 h after emergence [18]. It is unknown, however, whether this decline leads to increased de novo production of fatty acids and whether lipogenesis is restricted to the production of free fatty acids or also comprises TAGs for energy storage. Remarkably, the male sex pheromone in *U. rufipes* is synthesized independently from fatty acid metabolism [35]. This demonstrates that lipogenesis is not restricted to species that use fatty acid-derived pheromones and raises the question whether lipogenesis is common in parasitic wasps rather than missing in most species.

In the present study, we investigated eleven additional species of parasitoid wasps from six families with respect to their ability to synthesize fatty acids de novo from carbohydrates. We performed feeding experiments with fully ^13^C-labeled α-(+)-D-glucose and analysed the insect-derived fatty acid methyl esters (FAME) by GC/MS for the incorporation of sugar-derived carbon. Using the model organism Nv, we furthermore investigated whether ebbing lipid resources in females due to increasing age and egg laying activity result in increased lipogenesis and whether de novo synthesized fatty acids are incorporated into TAGs. Our results demonstrate that all species investigated are capable of synthesizing fatty acids de novo from α-(+)-D-glucose and suggest emphatically that the presumed ‘lack of lipogenesis’ in parasitoid wasps needs to be re-evaluated.

## Results

### ^13^C-labeling experiments with eleven parasitoid species

We offered females of *Dibrachys cavus* (Dc)*, Muscidifurax raptorellus* (Mr)*, M. uniraptor* (Mu), *Lariophagus distinguendus* (Ld) (Pteromalidae), *Tachinaephagus zealandicus* (Tz), *Exoristobia phillipinensis* (Ep) (Encyrtidae), *Baryscapus tineivorus* (Bt) (Tineidae), *Cephalonomia tarsalis* (Ct) (Betylidae), *Trichogramma evanescens* (Trichgrammatidae), and *Habrobracon hebetor* (Hh) (Braconidae) a 10% solution of fully ^13^C-labeled α-(+)-D-glucose for two days. Unfed females served as controls. After this period, females were frozen and homogenized in groups of 5 (Hh: 2, Ct: 10, Te: 50-100) individuals in dichloromethane to extract the raw lipids (n=3 per species and treatment). Fatty acid derivatives were transesterified with acetyl chloride/methanol and extracted FAME were analysed by GC/MS. To infer ^13^C-incorporation into fatty acids, we focused on the diagnostic ion pair m/z 90 (^13^C-labeled) and m/z 87 (unlabelled) in the mass spectra of palmitic acid methyl ester (PAME) and stearic acid methyl ester (SAME). We concluded ^13^C-incorporation into PAME and SAME from (a) a calculated incorporation rate of >0.05% of the diagnostic ion m/z 90 (peak area m/z 90 in relation the peak areas of m/z 90 + m/z 87) and (b) a slightly reduced retention time (ca. 1.5 s) of m/z 90 compared to m/z 87 (Fig. S1 in the supplementary material) [28, 36]. Female-derived chromatograms of all eleven species showed the diagnostic ion m/z 90 at the expected retention times of PAME and SAME (Figs. 1a-l,a-j, Tab. S1 in the supplementary material) demonstrating that all of them are able to synthesize fatty acids de novo from glucose.

**Fig. 1.**
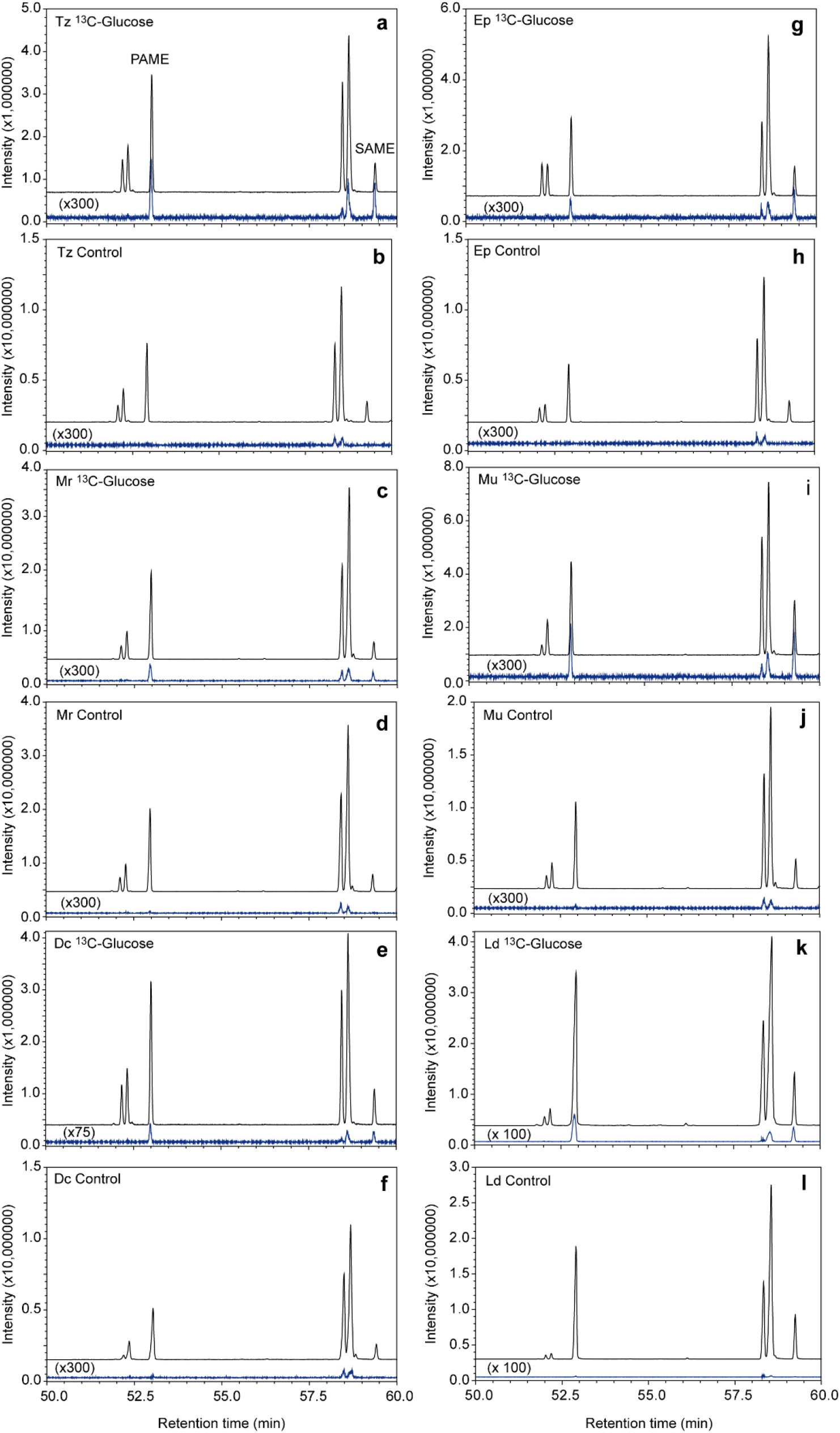
GC/MS analysis of transesterified lipid extracts from female parasitoids I. (**a**) *Tachinaephagus zealandicus* (*Tz*), (**c**) *Muscidifurax raptorellus* (Mr), (**e**) *Dibrachys cavus* (Dc), (**g**) *Exoristobia phillipinensis* (Ep), (**i**) *Muscidifurax uniraptor* (Mu), and (**k**) *Lariophagus distinguendus* (Ld) wasps fed fully ^13^C-labeled α-D-glucose for two days. Panels **b**, **d**, **f**, **h**, **j**, and **l** show the chromatograms of the respective unfed control wasps. Each panel shows the total ion chromatograms (upper trace) and the extracted ion chromatogram of the diagnostic ion m/z 90 (blue lower trace, magnification factors given in brackets). The peaks of palmitic acid methyl ester (PAME) and stearic acid methyl ester (SAME) are indicated in panel **a**.

Extracted ion chromatograms of the diagnostic ion m/z 90 on a 60 m high-resolution GC-column showed the expected retention time shift of 1.5 s in the samples from glucose-fed wasps due to the inverse isotope effect of heavier isotopes [36] (Fig. 3a-b). Across all species, the calculated incorporation rates of wasps having fed on ^13^C-labeled glucose were significantly higher (mean ± SEM, PAME: 1.8 ± 0.44%; SAME: 2.3 ± 0.4%) than in the unfed controls (PAME: 0.03 ± 0.003; SAME: 0.02 ± 0.004) (t-test: p < 0.0001 for both PAME and SAME). Some samples of ^13^C-glucose-fed females with higher calculated ^13^C-incorporation rates (>0.5%) showed additionally increased abundances of the diagnostic ion at m/z 150 (unlabelled: m/z 143) and even the fully labeled molecular ions (PAME m/z 286; SAME m/z 316; unlabelled: PAME m/z 270, SAME: 298) (Figs. S2c and S3b in the supplementary material). Magnification of the molecular ion region of the mass spectra furthermore revealed the presence of clusters of partially ^13^C-labeled molecular ions (PAME: m/z 280, 282, 284; SAME: m/z 308, 310, 312, 314) resulting from the incorporation of a varying number of ^13^C-labeled acetate units. In samples from Ld, Ac, Ct, Hh, Te, and Nv, the respective isotope clusters were also detectable in the methyl esters of the unsaturated fatty acids oleic acid and linoleic acid (Figs. S4, S5 in the supplementary material) demonstrating that ^13^C-labeled stearic acid had been processed by Δ9- and Δ12-desaturares [37].

### Impact of age and oviposition on lipogenesis

To investigate whether parasitoids invest increasingly in lipogenesis when their lipid reserves get depleted, we performed a feeding experiments with Nv. We offered ^13^C-labeled α-D-glucose for 2d to females that differed in age and oviposition history. For this purpose, we provided newly emerged females for 0, 2, or 4d with hosts ad libitum prior to glucose feeding. We expected a depletion of lipid reserves with increasing age and oviposition history. Newly emerged females without prior access to hosts and glucose served as control. After the feeding period, we extracted the raw lipids from individual females and analysed by GC/MS the total amount of FAME as well as the ^13^C-incorporation rates into PAME and SAME. The total amounts of FAME in newly emerged females having access to glucose for 2d did not differ from the lipid levels of the control wasps while females with a 2-d or 4-d oviposition history contained significantly lower amounts of fatty acid derivatives (Fig. 4a, Tab. S3). Lipogenesis, however, was detectable in females from all three treatment groups irrespective of whether they were lipid-limited or not (Fig. 4b-c). Additionally, the calculated relative ^13^C-incorporation rate into PAME was significantly higher in females with a 4-d oviposition history when compared to those that were not given the opportunity to lay eggs (Fig. 4b).

### De novo biosynthesis of TAGs

Finally, we investigated in Nv whether de novo synthesized fatty acids are incorporated into TAGs. For this purpose, we extracted pools of three Nv females of differing age and oviposition history from the ^13^C feeding experiment (see above) and fractionated the resulting raw lipids by size exclusion high performance liquid chromatography (HPLC) (n=3 per treatment). This technique allows for the isolation of TAGs as a single chromatographic peak [38] (Fig. S6 in the supplementary material). We transesterified the isolated TAGs from the differently treated females and determined the total amounts of FAME as well as the ^13^C incorporation into PAME and SAME. ^13^C-labeled PAME and SAME were detectable in the TAGs of ^13^C-glucose-fed Nv females irrespective of their age/oviposition history at calculated incorporation rates of up to 28% (PAME) and 49% (SAME), respectively (Fig. 2k-l, Tab. S2).

**Fig. 2.**
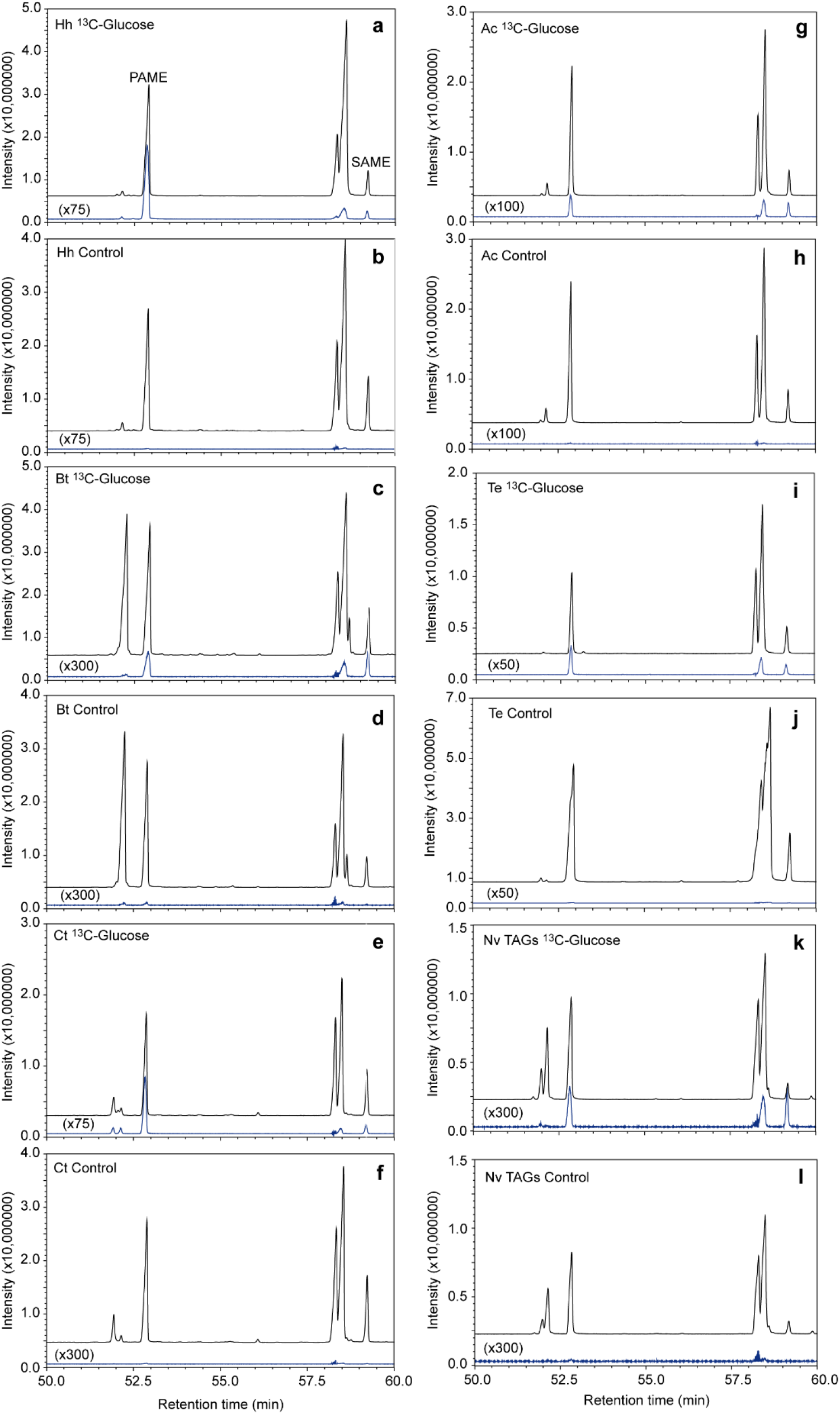
GC/MS analysis of transesterified lipid extracts from female parasitoids II. (**a**) *Habrobracon hebetor* (Hh), (**c**) *Baryscapus tineivorus* (Bt), (**e**) *Cephalonomia tarsalis* (Ct), (**g**) *Anisopteromalus calandrae* (Ac), (**i**) *Trichogramma evanescens* (Te), and (**k**) *Nasonia vitripennis* (Nv) fed fully ^13^C-labeled α-D-glucose for two days. Panels **b**, **d**, **f**, **h**, **j**, and **l** show the chromatograms of the respective unfed control wasps. Lipids of Nv were fractionated prior to transesterification by size exclusion HPLC (see Fig. S6 in the supplementary material) and represent the triacylglyceride fraction (TAGs). Each panel shows the total ion chromatograms (upper trace) and the extracted ion chromatogram of the diagnostic ion m/z 90 (lower trace, magnification factors given in brackets). The peaks of palmitic acid methyl ester (PAME) and stearic acid methyl ester (SAME) are indicated in panel **a**.

**Fig. 3.**
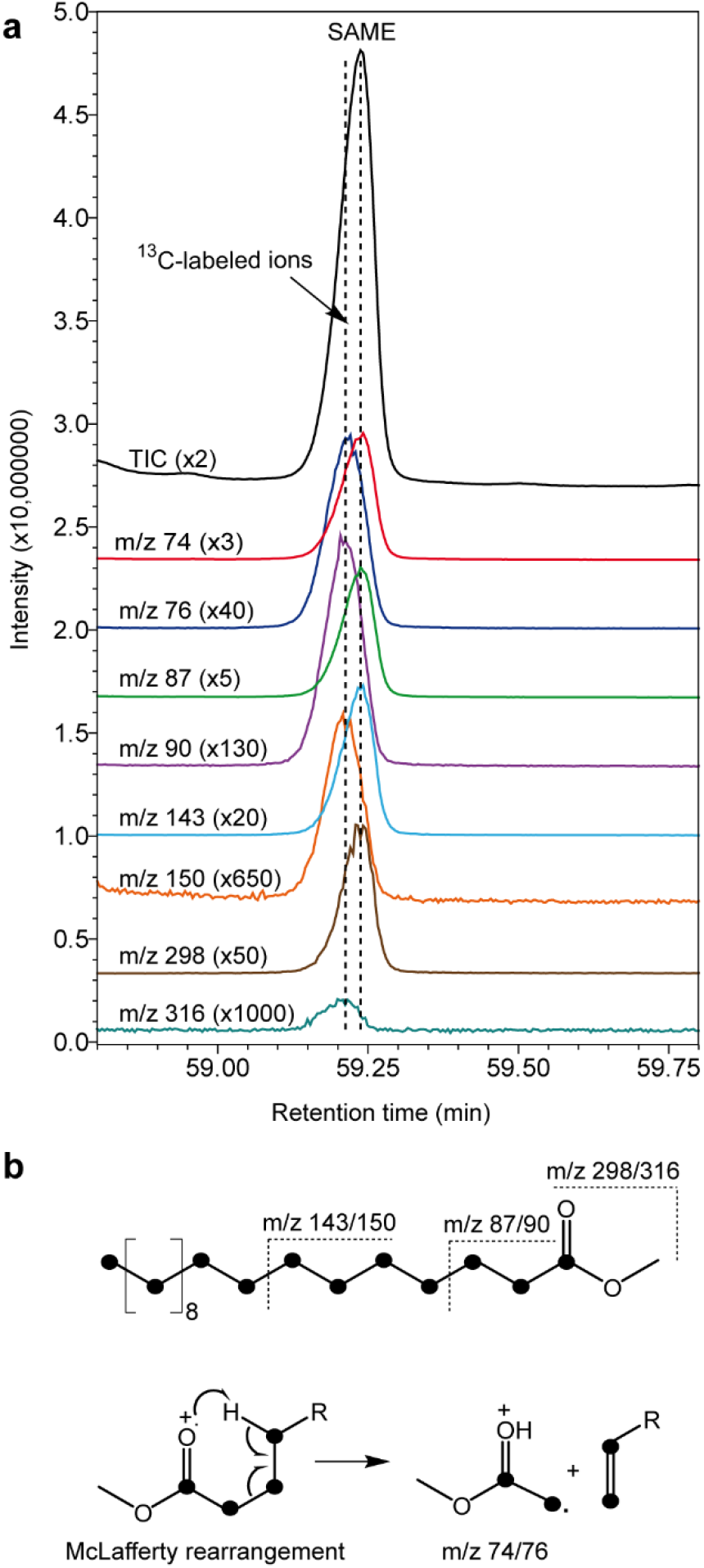
Diagnostic ions for the detection of ^13^C-incorporation into saturated fatty acids. (**a**) Elution profile of stearic acid methyl ester (SAME) from *Habrobracon hebetor* females fed fully ^13^C-labeled α-D-glucose. Shown are total ion current chromatogram (TIC) and extracted ion chromatograms of the diagnostic ion pairs (unlabelled/fully ^13^C-labeled) m/z (74/76), (87/90), (143/150), and (298/316), magnification factors are given in brackets). Dotted lines indicate the peak maxima of labelled and unlabelled diagnostic ions to demonstrate the slightly decreased retention time of the labelled ions due to the inverse isotope effect of heavier isotopes. (**b**) Structures of the mass spectrometric diagnostic ion pairs monitored in panel (**a**). Black dots represent ^13^C atoms in the labelled ions.

**Fig. 4.**
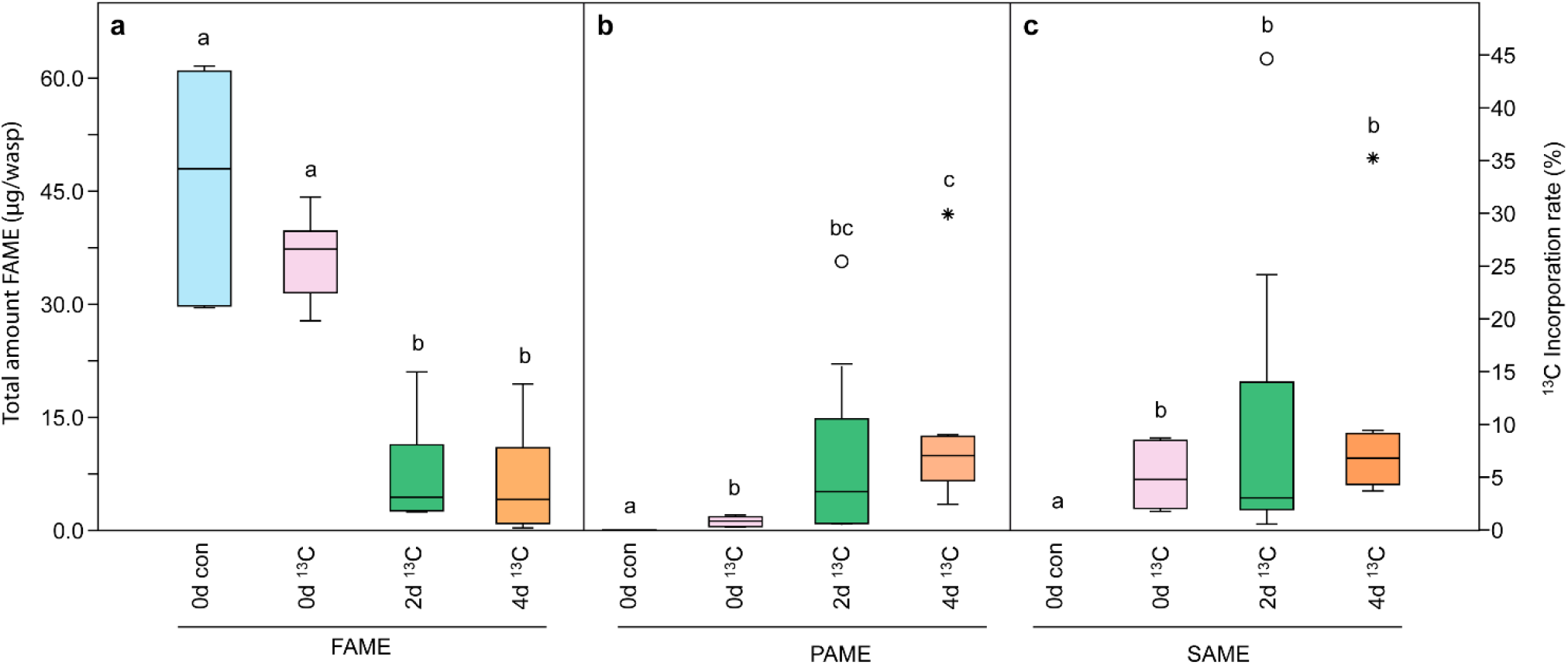
Effect of age and oviposition history on lipogenesis in *Nasonia vitripennis.* (**a**) Total amounts of fatty acid methyl esters (FAME) and (**b**-**c**) calculated ^13^C-incorporation rates into palmitic acid methyl ester (PAME) and stearic acid methyl ester (SAME) from transesterified total lipids of female *Nasonia vitripennis* females. Females had either 0 (newly emerged), 2, or 4 days ad libitum access to hosts and were then fed fully ^13^C-labeled α-D-glucose (0d^13^C, 2d^13^C, 4d^13^C) for two days. Newly emerged females without access to ^13^C-labeled α-D-glucose were used as control (0d con). Different lowercase letters within each panel indicate significant differences at p<0.05 (Kruskal-Wallis test followed by Bonferroni-corrected multiple Mann-Whitney U-tests, n=6).

## Discussion

The present study clearly demonstrates that all investigated parasitoid species belonging to six families are capable of synthesizing fatty acids de novo from α-D-glucose. Together with the four species of the *Nasonia* group reported earlier [28], we have now proven de novo biosynthesis of fatty acids in 15 species and so far we did not study a single species in which fatty acid biosynthesis was undetectable. Our experiments with Nv demonstrated furthermore that even newly emerged parasitoids with full lipid reserves start to synthesize fatty acids upon sugar feeding and convert them, at least partly, into TAGs. We therefore conclude that the presumed ‘lack of lipogenesis’ in parasitoid wasps [8, 9] needs to be re-evaluated. We propose that lipogenesis in parasitoids is the rule rather than the exception. Parasitoid wasps, like most organisms on earth [7], possess and use the enzymatic machinery to convert glucose into fatty acids and their derivatives. The total mass of fatty acids declined drastically in Nv females after 2-4 days with the opportunity to lay eggs. With declining lipid reserves, the calculated relative ^13^C-incorporation rates increased in PAME and remained constant in SAME reaching levels up to 28% and 49%, respectively. In several but not all species studied so far with respect to lipogenesis, total lipid mass declined although a carbohydrate source was offered ad libitum [8, 9, 13–18]. In the light of the present study, this can be explained by lipogenesis only partially compensating the depletion of the lipid reserves. Lipid reserves of parasitic wasps may deplete quickly particularly in females due the energy demands during host finding and oviposition [18]. Approximately 30-40% of the dry weight of insect oocytes consists of lipids, the vast majority of which are provided by the mother [39]. The fact that lipogenesis can occur in parasitoids despite a drastic decline of total lipids suggests that studies having relied on bulk methods for lipid quantification have overlooked the ability of parasitoids to synthesize fatty acids de novo. Visser et al. [9], for instance, fed 21 parasitoid species honey ad libitum and found five of them gaining and 11 losing lipid mass, while five showed stable levels. The present study demonstrates that a ‘lack of lipogenesis’ in species with declining or stable lipid levels despite having access to carbohydrates should no longer be claimed unless being verified by stable isotope labelling techniques. Not all of these labelling techniques, however, appear to be suitable to demonstrate lipogenesis in parasitoid wasps. A recent study, for instance, involving feeding experiments with unlabelled glucose dissolved in deuterated water, failed to detect lipogenesis in *N. vitripennis* but worked well with honeybees [40]. Unlike the method reported here, this method does not monitor the direct incorporation of sugar-derived atoms into fatty acids but the indirect incorporation of deuterium during fatty acid biosynthesis via deuterated NADPH after H-D exchange [41]. The labeling and fractionation techniques reported here allow for the sensitive detection of lipogenesis even in single specimens of these mostly tiny species and thus offer excellent opportunities for the in-depth study of more taxa as well as the biotic and abiotic factors controlling the degree at which lipogenesis occurs in parasitoid wasps.

## Materials and Methods

### Insects

Dc, Nv (both originally collected from empty bird’s nests near Hamburg in Northern Germany), Mr, Mu (kindly provided by E. Verhulst, University of Wageningen), Tz (collected from carcass baits near Gießen, Germany), and Ep (collected from carcass baits in Davie, Florida, USA) were reared on juvenile stages of the green bottle fly *Lucilia caesar* (Diptera: Calliphoridae). Dc, Nv, Mr, and Mu were reared on pupae that were freeze-killed 2d after pupation and thawed on demand [42]. After thawing and drying the hosts for 2 h at 30 °C, these were exposed in Petri dishes to newly emerged wasps and kept in an incubator at 25 °C and 50% relative humidity. Tz and Ep females were provided with live final-instar larvae (Tz) or live pupae (0-2 days after pupation, Ep), and thereafter reared at identical conditions as mentioned above. A starter culture of Ld was obtained from a local company (Biologische Beratung Prozell und Schöller GmbH, Berlin, Germany) and reared on larvae of the granary weevil *Sitophilus granarius* (Coleoptera: Curculionidae) as described previously [43]. Bt, reared on larvae of the clothing moth *Tineola bisselliella* (Lepidoptera: Tineidae), Ct, reared on larvae of the sawtooth grain beetle *Oryzaephilus surinamensis* (Coleoptera: Silvanidae), Ac (reared larvae of *S. granarius*), Te, reared on eggs of the Mediterranean flour moth *Ephestia kuehniella* (Lepidoptera: Pyralidae), and Hh, reared on larvae *E. kuehniella*, were also obtained from Biologische Beratung Prozell und Schöller GmbH, Berlin, Germany and used as delivered.

### Feeding experiments

Newly emerged, mated females of Dc, Ld, Mr, Mu, Tz, and Ep were provided with suitable hosts ad libitum for oviposition for two days to ensure that they could metabolize parts of their existing fat reserves. After this pre-treatment, females were kept for two more days in groups of 5 wasps (n=3 for each species) in 1.5 ml microcentrifuge tubes, the bottom of which was covered by 40 μl of a 10% solution of fully ^13^C-labeled α-D-(+)-glucose (99% ^13^C, Sigma-Aldrich, Taufkirchen, Germany). The labelled glucose solution was renewed after 24 h when necessary. Control wasps were kept under identical conditions without providing glucose solution. Females of Ac, Ct, Bt, Te, and Hh were used without prior pre-treatment as delivered by the supplier. After the feeding period, wasps were frozen and kept at −20 °C until being used for chemical analyses. To investigate the influence of age and oviposition on lipogenesis in more detail, we performed another feeding experiment with Nv. Prior to offering the ^13^C-labeled α-D-(+)-glucose solution for two days, newly emerged females were exposed for 0 (no hosts offered), 2, or 4 days to hosts ad libitum. For control, newly emerged females were frozen on the day of emergence without prior oviposition and sugar feeding (n=6 per treatment). These wasps were analysed singly by GC/MS to determine total amounts of fatty acids and ^13^C incorporation rates into PAME and SAME as described below. Furthermore, we used Nv females from this feeding experiment to isolate the TAGs by size exclusion HPLC as described below.

### Isolation of TAGs by size exclusion HPLC

Pools of three Nv females per sample and treatment from the ^13^C-glucose feeding experiment (n=3 for each treatment) were homogenized with 200 μl dichloromethane. After 30 min, the extracts were transferred to a glass vial with a 300 μl glass insert and concentrated under a stream of nitrogen to 20 μl. These sample were completely analysed by size exclusion HPLC using a method described previously [38]. Analyses were performed on an Agilent Infinity 1260 HPLC system equipped with a 300 × 7.5 mm PLgel SE-HPLC column (particle size 5 μm, pore size 100 Å) (Agilent Technologies, Waldbronn, Germany). Dichloromethane was used as eluent at a flow rate of 1.00 ml/min. Eluting analytes were detected with an Infinity 1260 multi-wavelength UV/VIS detector at 254 nm. Under these conditions, TAGs elute as a single peak between 5.8 and 6.5 minutes [38]. To confirm this elution range, we analysed 20 μl of a dichloromethane solution (1 μg/μl) of synthetic triolein ((*Z*)*-*9-octadecenoic acid 1,2,3-propanetriyl ester, Sigma-Aldrich, Deisenhofen, Germany). The TAG fractions of all samples were collected with an Agilent Infinity 1260 fraction collector. After adding 10 μl of a dichloromethane solution of tetracosane (C24, 50 ng/μl), fractions were carefully dried under a stream of nitrogen before being used for transesterification (see below).

### Transesterification of lipids for GC/MS analysis

After thawing, groups of five (*Hh*: 2, Bt: 10) females or 50-100 Te of both sexes (n = 3 for each species and treatment) were homogenized with a glass pestle in a 1.5 ml glass vials after adding 200 μl dichloromethane. The homogenates were kept at room temperature for 30 min. Subsequently, the extracts were transferred to new 1.5 ml glass vials. The residues were washed with additional 200 μl dichloromethane and the unified extracts were carefully dried under a stream of nitrogen. The residues were re-suspended in 200 μl of methanol and 20 μl of acetyl chloride (10%, dissolved in methanol) and transesterified for 3 h at 80°C. Thereafter, 200 μl of a solution of sodium hydrogen carbonate (5%) were added and FAME were extracted with 200 μl hexane. The hexane phase was concentrated to 25 μl under nitrogen and used for GC/MS analysis. The same protocol was used for transesterification of the TAG fractions from Nv females. Sample preparation of Nv females from the ^13^C glucose feeding experiment was slightly modified. Here, females of equal size were extracted singly (n = 6 for each of the four treatments) with 100 μl dichloromethane containing 2 ng/μl C24 as an internal standard and washed with another 100 μl dichloromethane.

### Chemical analysis by GC/MS

Chemical analyses were performed using a Shimadzu QP2010 Plus GC/MS system equipped with a 60 m x 0.25 mm inner diameter BPX5 capillary column (film thickness 0.25 μm, SGE Analytical Science Europe, Milton Keynes, UK). Samples (1 μl, Nv and Te: 2 μl) were injected splitless at 300°C using a Shimadzu AOC 20i auto sampler. The MS was operated in the electron impact ionization mode at 70 eV; the mass range was m/z 35-500. Helium was used as carrier gas at a constant velocity of 40 cm s^−1^. The temperature program started at 50°C, increased at 3°C min^−1^ to 280°C and was kept at this temperature for 22 min. Identification of FAME was done by comparison of retention times and mass spectra with those of authentic reference chemicals (FAME: reference mixture of 37 FAME, Sigma-Aldrich, Deisenhofen, Germany). As labelled reference, we analysed fully ^13^C-labeled SAME (Sigma-Aldrich) and a mixture of ^13^C-labeled and unlabelled SAME that were esterified with unlabelled acetyl chloride/methanol. For the detection of incorporated ^13^C in insect-derived fatty acids we focused on fully ^13^C-labeled palmitic acid methyl ester (PAME) and stearic acid methyl ester (SAME). Mass spectra of these compounds are less complex than those of unsaturated FAME facilitating the interpretation of the results. To infer ^13^C-incorparation into saturated fatty acids of all samples, we monitored the diagnostic ion m/z 90 (unlabelled: m/z 87, see Fig. 2H) which occurs in mass spectra of both ^13^C-labeled PAME and SAME [28]. For selected samples with high ^13^C-incorporation (>0.5%), we also monitored the diagnostic ions m/z 150 (unlabelled ion: m/z 143) and the molecular ions m/z 286 (PAME) and 316 (SAME, unlabelled ions: m/z 270 and 298, respectively). On the high-performance 60 m capillary GC-column used in this study, ^13^C-incorparation of labelled ions is furthermore supported by a slightly decreased (ca. 1.5 s) retention time compared to the unlabelled analogues due to the inverse isotope effect of heavier isotopes (Fig. S1) [36]. Labelling rates in PAME and SAME were calculated by relating the peak area of the labelled diagnostic ion m/z 90 to the added peak areas of the respective unlabelled (m/z 87) and labelled diagnostic ions [28]. At high injected amounts, mass spectra of unlabelled PAME and SAME may show a weak signal at m/z 90, which however never exceeds 0.05% of the added peak area of m/z 90 + m/z 87 (see Tabs. S1, S2 in the supplementary material) and does not show the shifted retention time in the extracted ion chromatogram. As the structure of the diagnostic ion m/z 90 reveals (see Fig. 3b), the calculation of ^13^C-incorporation is very conservative, because it considers only those fatty acid molecules with the first two C_2_ units originating from ^13^C-glucose-derived acetyl CoA, while those fatty acid molecules carrying labelled C_2_-units exclusively at the subsequent positions of the fatty acid chain are not included. Hence, the calculated incorporation rates can be used for comparisons between treatments but do not allow conclusions about the exact proportion of fatty acids having been synthesized from the ingested ^13^C-labeled glucose. For quantification of FAME in the Nv samples we integrated the peak areas of the six most abundant FAME (C16:0, C16:1^Δ9^, C16:1^Δ7^, C18:0, C18:1^Δ9^, and C18:2^Δ9,12^) and related them to the peak area of the internal standard C24.

### Statistical analysis

Statistical analyses were performed with PAST 4.03 scientific software [44]. Calculated ^13^C-incorporation rates in PAME and SAME from ^13^C-glucose fed and unfed wasps were compared across species with a t-test for unpaired samples. Total FAME amounts and calculated ^13^C incorporation rates in PAME and SAME in the lipid extracts from Nv females of different age and oviposition history were compared by a Kruskal-Wallis H-test followed by Bonferroni-corrected multiple Mann-Whitney U-tests.

## Acknowledgements

This paper is dedicated to Prof. Dr. Dr. h.c. mult. Wittko Francke who recently passed away much too early. The authors thank Sonja Fleischmann and Sophia Goedecke for rearing the insects.

## Author Contributions

JR designed the experiments, developed the methods, performed parts of the experiments, analysed the results, and drafted the manuscript. LP performed parts of the experiments and analysed the data. TP provided two species of the studied parasitic wasps and revised the manuscript.

