## supplementary material for "Parasitic wasps do not lack lipogenesis"

**Electronic supplementary material**

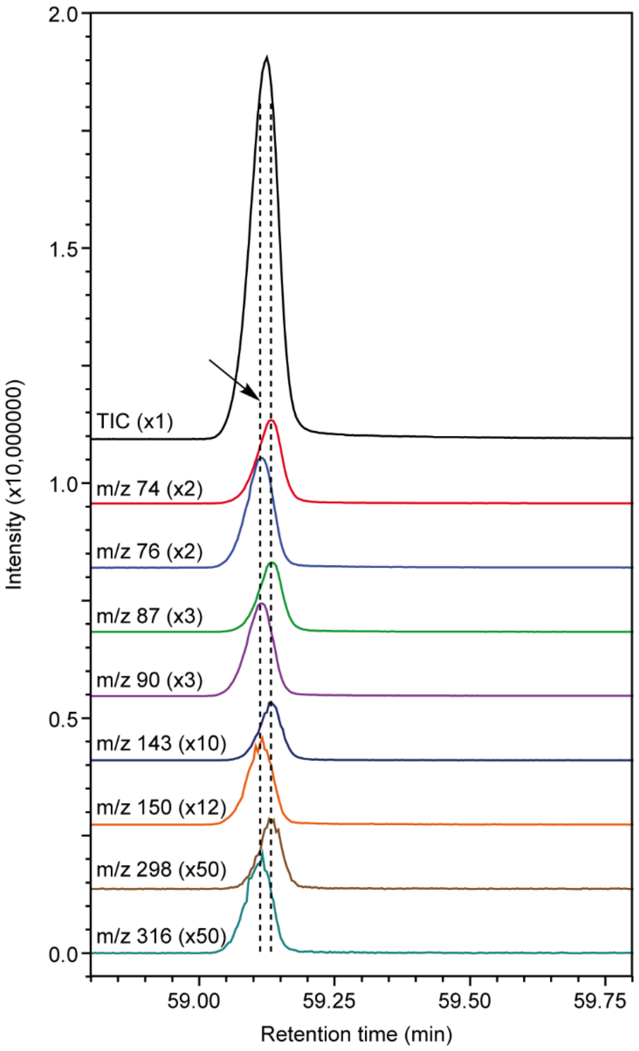


**Figure S1. Elution pattern of selected $^{13}C$-labeled and unlabeled diagnostic ions from synthetic stearic acid methyl ester (SAME).** Elution profile of a synthetic blend of fully $^{13}C$-labeled and unlabeled SAME. Shown are total ion current chromatogram (TIC) and extracted ion chromatograms of the diagnostic ion pairs (unlabeled/fully $^{13}C$-labeled) m/z (74/76), (87/90), (143/150), and (298/316), magnification factors are given in brackets). Dotted lines indicate the peak maxima of labeled and unlabeled diagnostic ions to demonstrate the slightly decreased (1.5 s) retention time of the labeled ions due to the inverse isotope effect of heavier isotopes. Structures of the mass spectrometric diagnostic ion pairs are given in Figure 3b of the main paper.

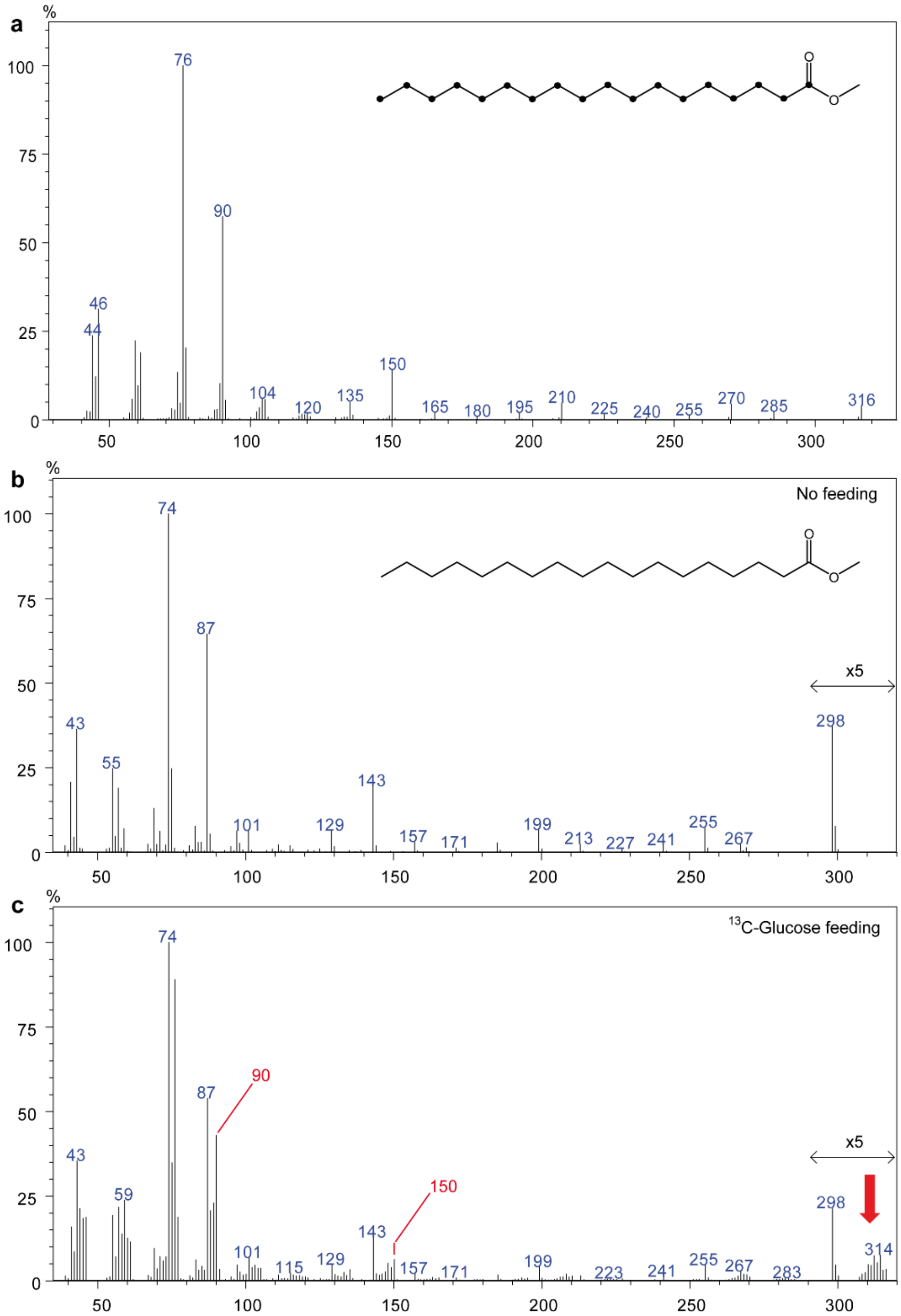


**Figure S2. $^{13}$C-incorporation into stearic acid methyl ester.** Mass spectra of (**a**) fully $^{13}$C-labeled stearic acid methyl ester (SAME), and SAME obtained by transesterification of lipid extracts from (**b**) unfed *Habrobracon hebetor* (Hh) females or (**c**) Hh females fed fully $^{13}$C-labeled α-D-glucose for 2 days. Diagnostic ions indicating the incorporation of $^{13}$C in panel (C) are indicated in red (for structures of the ions see Fig. 3b of the main paper). The red arrow shows the magnified region next to the molecular ion where a cluster of fully (m/z 316) and partially labeled molecular ions can be seen which result from the incorporation of a varying number of $^{13}$C-labeled acetate units into the fatty acid chain.

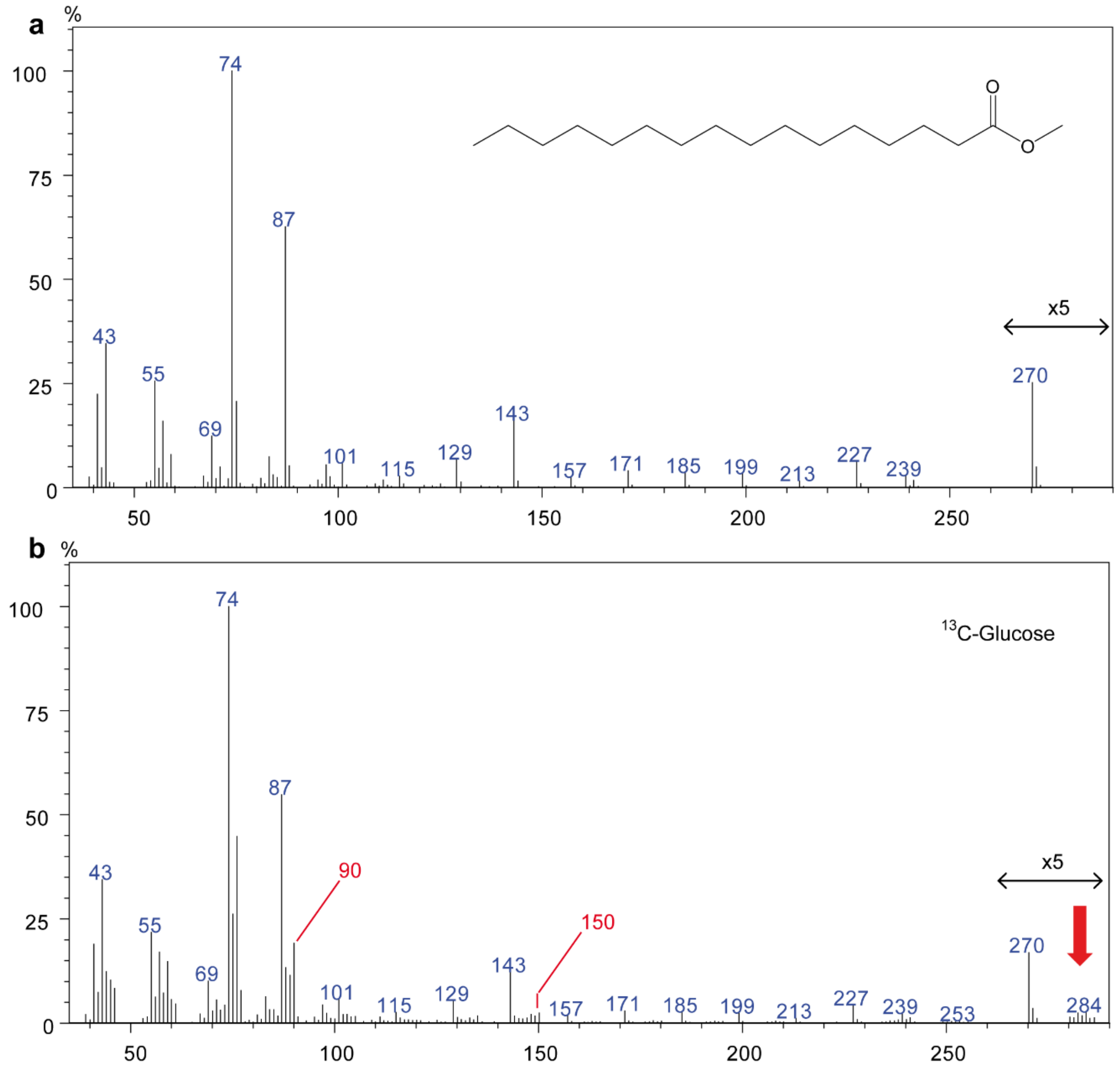


**Figure S3. $^{13}$C-incorporation into palmitic acid methyl ester.** Mass spectra of palmitic acid methyl ester obtained by transesterification of lipid extracts from (**a**) unfed *Habrobracon hebetor* (Hh) females or (**b**) Hh females fed fully $^{13}$C-labeled α-D-glucose for 2 days. Diagnostic ions indicating the incorporation of $^{13}$C in the bottom spectrum are indicated in red (for structures of the ions see Fig. 3b of the main paper). The red arrow shows the magnified region next to the molecular ion where a cluster of fully (m/z 286) and partially labeled molecular ions can be seen which result from the incorporation of a varying number of $^{13}$C-labeled acetate units into the fatty acid chain.

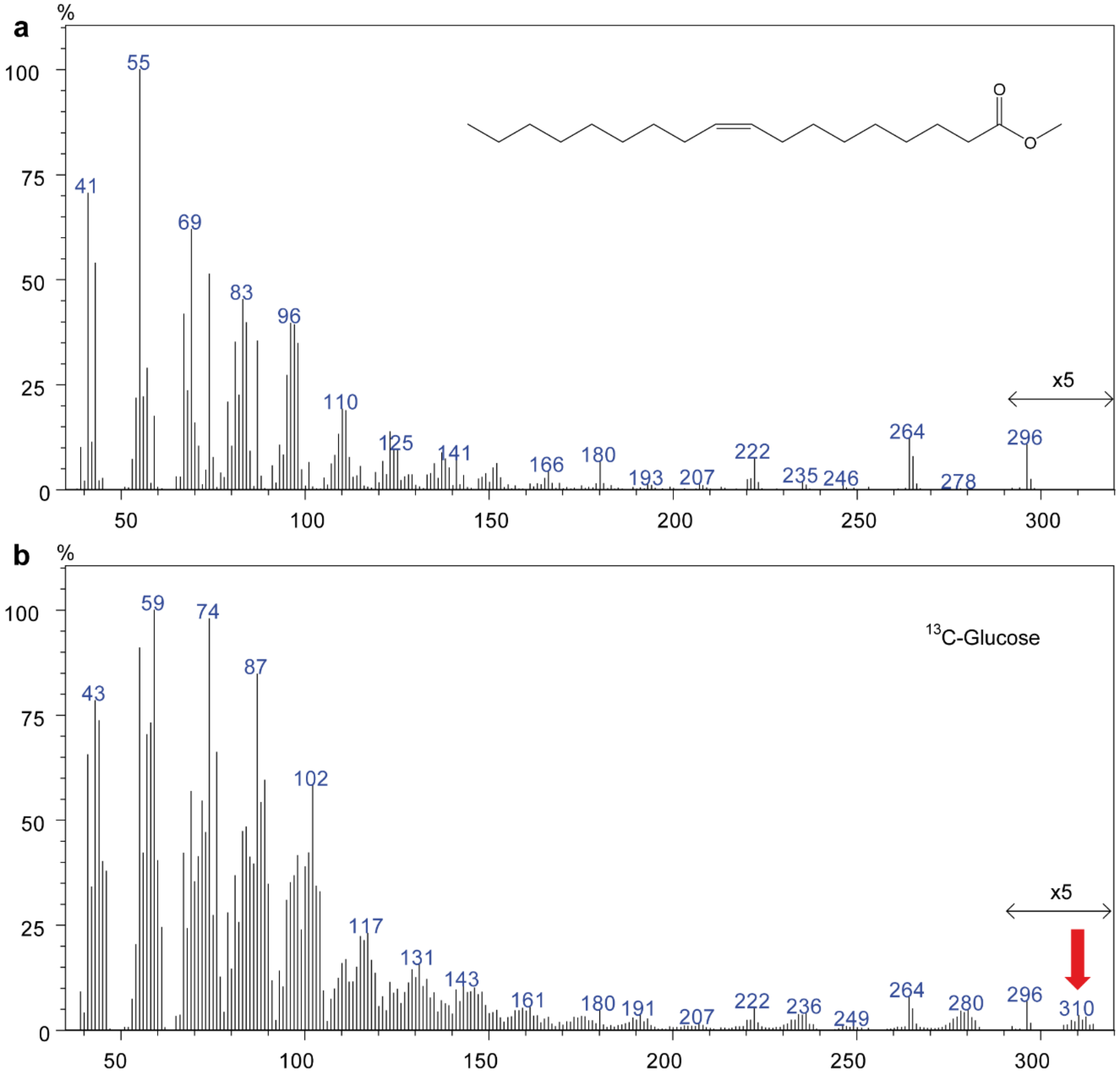


**Figure S4. $^{13}$C-incorporation into oleic acid methyl ester.** Mass spectra of oleic acid methyl ester obtained by transesterification of lipid extracts from (**a**) unfed *Nasonia vitripennis* females or (**b**) females fed fully $^{13}$C-labeled α-D-glucose for 2 days. The red arrow shows the magnified region next to the molecular ion where a cluster of fully (m/z 314) and partially labeled molecular ions can be seen which result from the incorporation of a varying number of $^{13}$C-labeled acetate units into the fatty acid chain.

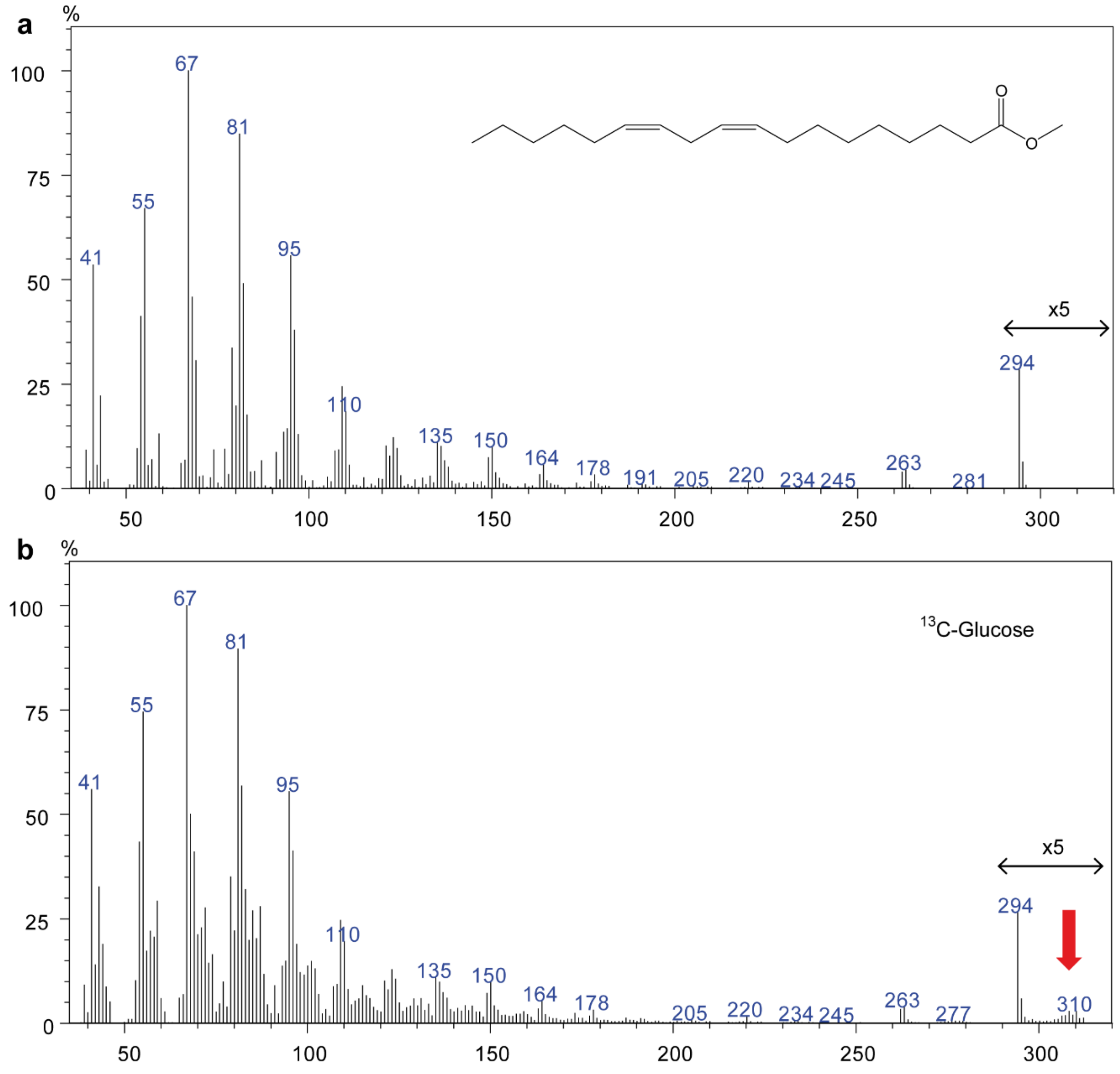


**Figure S5. $^{13}$C-incorporation into linoleic acid methyl ester.** Mass spectra of linoleic acid methyl ester obtained by transesterification of lipid extracts from (**a**) unfed *Nasonia vitripennis* females or (**b**) females fed fully $^{13}$C-labeled α-D-glucose for 2 days. The red arrow shows the magnified region next to the molecular ion where a cluster of fully (m/z 312) and partially labeled molecular ions can be seen which result from the incorporation of a varying number of $^{13}$C-labeled acetate units into the fatty acid chain.

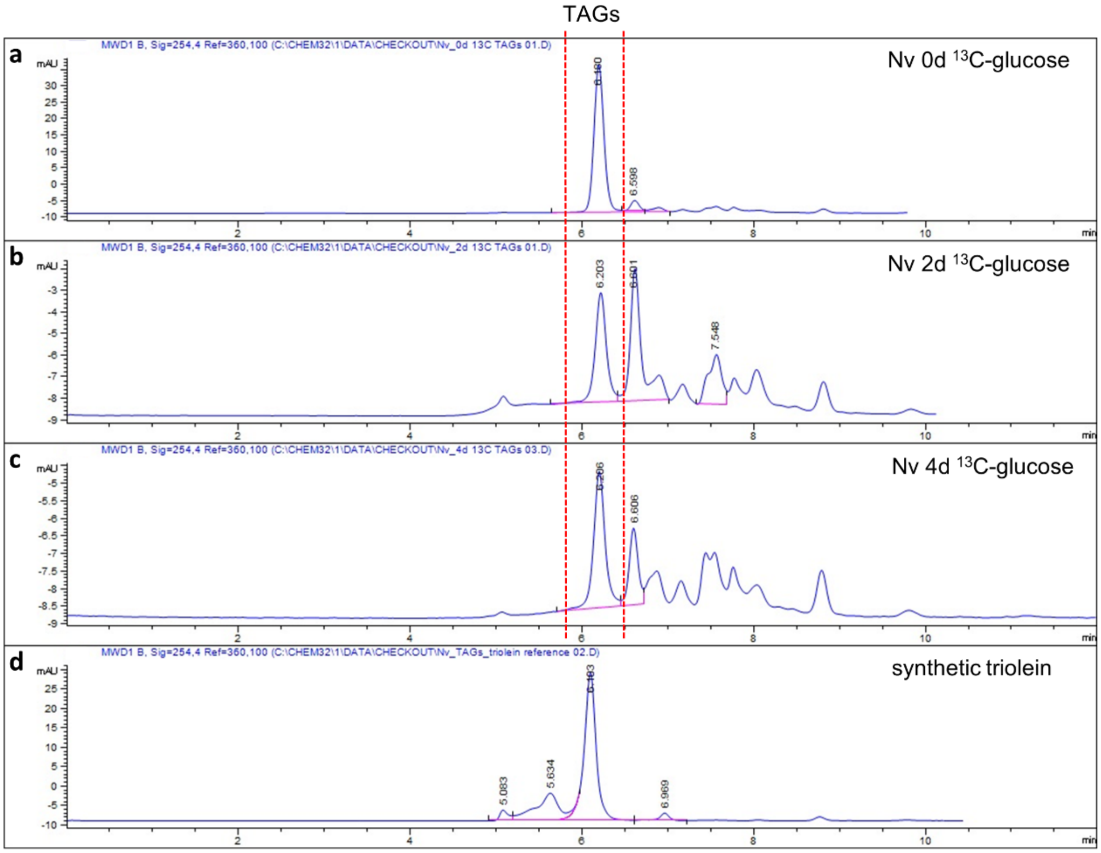


**Fig. S6. Isolation of triacylglycerides (TAGs) from *Nasonia vitripennis.*** Separation of raw lipid extracts (**a**-**c**) from *N. vitripennis.* Females were either newly emerged (0d), 2 or 4 days old (2d, 4d, with ad libitum access to hosts) and were fed fully $^{13}$C-labeled $\alpha$-D-glucose for two additional days and (**d**) synthetic triolein on a 300 × 7.5 mm PLgel size exclusion HPLC column (particle size 5 µm, pore size 100 Å) operated with dichloromethane as a mobile phase at a flow rate of 1 ml/min. Elution profiles were detected by a UV detector at 254 nm. Three wasps were pooled per sample for lipid extraction. Dotted lines indicate the fractions eluting between 6.8–7.5 min containing the TAGs which were collected for GC/MS analysis of fatty acid methyl esters.

**Tab. S1.** Calculated incorporation rates of $^{13}C$ into palmitic acid methyl ester (PAME) and stearic acid methyl ester (SAME) from transesterified lipids of female parasitic wasps fed fully $^{13}C$-labeled $\alpha$-D-glucose. Incorporation was calculated by relating the peak area of the $^{13}C$-labeled diagnostic ion m/z 90 to the added peak areas of m/z 90 and the respective unlabelled ion m/z 87. Unfed wasps of either species were analyzed for control (Con). Asterisks indicate control samples in which m/z 90 was detected in traces. The respective peaks, however, did not show the typical decreased retention times (ca. 1.5 s) due to the inverse isotope effect. Calculated incorporation rates >0.05% indicate incorporation of $^{13}C$-labeld acetyl-CoA units. Respective samples are printed in bold. Species names as follows: *Anisopteromalus calandrae* (*Ac*), *Dibrachys cavus* (*Dc*), *Lariophagus distinguendus* (*Ld*), *Muscidifurax raptorellus* (*Mr*), *M. uniraptor* (*Mu*), *Tachinaephagus zealandicus* (*Tz*), *Exoristobia phillipinensis* (*Ep*), *Cephalonomia tarsalis* (*Ct*), *Barsiscapus tineivorus* (*Bt*), *Trichogramma evanescens* (*Te*), and *Habrobracon hebetor* (*Hh*).

| Family | Species | Replicate | PAME | | SAME | |
|---|---|---|---|---|---|---|
| | | | $^{13}C$-glu | Con | $^{13}C$-glu | Con |
| Pteromalidae | *Ac* | 1 | **1.50** | 0.04* | **5.01** | 0.04* |
| | | 2 | **0.30** | 0.04* | **0.88** | 0.04* |
| | | 3 | **0.47** | 0.04* | **1.96** | 0.00 |
| | *Dc* | 1 | **4.36** | 0.00 | **4.98** | 0.00 |
| | | 2 | **9.23** | 0.00 | **8.86** | 0.00 |
| | | 3 | **1.03** | 0.04* | **1.44** | 0.00 |
| | *Ld* | 1 | **1.51** | 0.03* | **2.39** | 0.04* |
| | | 2 | **0.12** | 0.04* | **0.71** | 0.05* |
| | | 3 | **0.44** | 0.04* | **0.62** | 0.01* |
| | *Mr* | 1 | **0.56** | 0.03* | **1.26** | 0.00 |
| | | 2 | **0.06** | 0.04* | **0.06** | 0.00 |
| | | 3 | 0.05 | 0.03* | 0.00 | 0.00 |
| | *Mu* | 1 | **0.30** | 0.00 | **0.84** | 0.00 |
| | | 2 | **0.26** | 0.04* | **0.83** | 0.00 |
| | | 3 | **0.33** | 0.00 | **0.90** | 0.00 |
| Encyrtidae | *Tz* | 1 | **4.06** | 0.00 | **6.09** | 0.00 |
| | | 2 | **0.53** | 0.00 | **1.47** | 0.00 |
| | | 3 | **1.23** | 0.04* | **3.04** | 0.00 |
| | *Ep* | 1 | **0.57** | 0.00 | **1.97** | 0.00 |
| | | 2 | **0.16** | 0.00 | **1.31** | 0.00 |
| | | 3 | **0.55** | 0.00 | **3.05** | 0.00 |
| Bethylidae | *Ct* | 1 | **1.39** | 0.04* | **0.51** | 0.05* |

| | | | | | | |
|---|---|---|---|---|---|---|
| | | 2 | **0.27** | 0.04* | **0.20** | 0.04* |
| | | 3 | **6.27** | 0.05* | **2.15** | 0.04* |
| Eulophidae | *Bt* | 1 | **0.11** | 0.05* | **0.09** | 0.00 |
| | | 2 | **0.06** | 0.03* | **0.07** | 0.00 |
| | | 3 | **0.56** | 0.04* | **1.55** | 0.03* |
| Trichogrammatidae | *Te* | 1 | **4.75** | 0.04* | **5.55** | 0.04* |
| | | 2 | **2.59** | 0.04* | **4.08** | 0.04* |
| | | 3 | **5.56** | 0.04* | **6.45** | 0.04* |
| Braconidae | *Hh* | 1 | **0.27** | 0.04* | **0.46** | 0.05* |
| | | 2 | **7.76** | 0.05* | **3.77** | 0.03* |
| | | 3 | **6.11** | 0.04* | **7.21** | 0.03* |

**Tab. S2.** Total amounts of triacylglycerides (TAGs) and calculated incorporation rates of $^{13}C$ into palmitic acid methyl ester (PAME) and stearic acid methyl ester (SAME) from transesterified TAGs of female *Nasonia vitripennis* females. Females had either 0, 2, or 4 days old ad libitum access to hosts and were fed fully $^{13}C$-labeled $\alpha$-D-glucose for two additional days. Three wasps of either treatment were pooled for each replicate. Prior to transesterification, TAGs of the individual samples were isolated by size exclusion HPLC. $^{13}C$-incorporation was calculated after GC/MS analysis by relating the peak area of the $^{13}C$-labeled diagnostic ion m/z 90 to the added peak areas of m/z 90 and the respective unlabelled ion m/z 87. Asterisks indicate control samples in which m/z 90 was detected in traces. The respective peaks, however, did not show the typical decreased retention times (ca. 1.5 s) due to the inverse isotope effect.

| Treatment | Replicate | TAGs (µg/sample) | $^{13}C$ incorporation (%) | |
|---|---|---|---|---|
| | | | PAME | SAME |
| 0d, no oviposition no feeding (control) | 1 | 29.5 | 0.03* | 0.00 |
| | 2 | 10.5 | 0.00 | 0.00 |
| | 3 | 47.1 | 0.04* | 0.00 |
| 0d, no oviposition 2d $^{13}C$-glucose | 1 | 49.2 | 1.04 | 6.6 |
| | 2 | 52.8 | 0.84 | 5.5 |
| | 3 | 41.4 | 0.71 | 4.9 |
| 2d, oviposition 2d $^{13}C$-glucose | 1 | 8.8 | 25.3 | 49.0 |
| | 2 | 11.5 | 2.6 | 4.3 |
| | 3 | 14.4 | 4.4 | 12.4 |
| 4d, oviposition 2d $^{13}C$-glucose | 1 | 1.1 | 5.2 | 5.0 |
| | 2 | 6.7 | 1.1 | 1.6 |
| | 3 | 2.6 | 28.0 | 37.8 |

**Tab. S3. Effect of age and oviposition history on lipogenesis in *Nasonia vitripennis.*** Statistical analysis of the total amounts of fatty acid methylesters (FAME) and calculated incorporation rates of $^{13}C$ into palmitic acid methyl ester (PAME) and stearic acid methyl ester (SAME) from transesterified total lipids of female *Nasonia vitripennis* females. Females had either 0, 2, or 4 days ad libitum access to hosts and were fed fully $^{13}C$-labeled α-D-glucose (0d$^{13}C$, 2d$^{13}C$, 4d$^{13}C$) for two additional days. Newly emerged females without access to $^{13}C$-labeled α-D-glucose were used as control (0d con).

Kruskal-Wallis H-test FAME

| | |
|---|---|
| H (chi2): | 19.07 |
| p (same): | **0.0002644** |

Bonferroni-corrected multiple Mann-Whitney U-tests

| | 0d con | 0d $^{13}C$ FAME | 2d $^{13}C$ FAME | 4d $^{13}C$ FAME |
|---|---|---|---|---|
| 0d con | | 1 | **0.03045** | **0.01449** |
| 0d $^{13}C$ FAME | | | **0.03045** | **0.01449** |
| 2d $^{13}C$ FAME | | | | 1 |
| 4d $^{13}C$ FAME | | | | |

Kruskal-Wallis H-test PAME

| | |
|---|---|
| H (chi2): | 19.41 |
| p (same): | **0.0002223** |

Bonferroni-corrected multiple Mann-Whitney U-tests

| | 0d con | 0d $^{13}C$ PAME | 2d $^{13}C$ PAME | 4d $^{13}C$ PAME |
|---|---|---|---|---|
| 0d con | | **0.02863** | **0.02863** | **0.01386** |
| 0d $^{13}C$ PAME | | | 0.7692 | **0,01449** |
| 2d $^{13}C$ PAME | | | | 0.8258 |
| 4d $^{13}C$ PAME | | | | |

Kruskal-Wallis H-test SAME

| | |
|---|---|
| H (chi2): | 15.42 |
| p (same): | **0.001364** |

Bonferroni-corrected multiple Mann-Whitney U-tests

| | 0d con | 0d $^{13}C$ SAME | 2d $^{13}C$ SAME | 4d $^{13}C$ FSME |
|---|---|---|---|---|
| 0d con | | **0.01667** | **0.01667** | **0.009541** |
| 0d $^{13}C$ SAME | | | 1 | 1 |
| 2d $^{13}C$ SAME | | | | 0.3673 |
| 4d $^{13}C$ SAME | | | | |
